## Supplementary Figure 1-18; Supplementary Table 1 for "MarsGT: Multi-omics analysis for rare population inference using single-cell graph transformer"

### MarsGT: Deep learning multi-omics analysis for rare populations from single cells

**Supplementary Fig. 1.** The detailed workflow of MarsGT for rare cell population identification.
**Supplementary Fig. 2.** Performance comparison of rare cell identification on 350 simulated datasets in terms of F1 score, Precision, and Recall.
**Supplementary Fig. 3.** Performance comparison of the major cell and rare cell population simultaneously identification ability on 350 simulated datasets in terms of NMI, Purity, and Entropy.
**Supplementary Fig. 4.** Performance comparison of the false positive rate on 50 simulated datasets in terms of F1 score, Precision, Recall, NMI, Purity, and Entropy.
**Supplementary Fig. 5.** Performance comparison of the different proportion rare cell population identification ability on 5 simulated datasets in terms of F1 score, Precision, Recall, NMI, Purity, and Entropy.
**Supplementary Fig. 6.** Robustness test. MarsGT runs 20 times on the independent test set.
**Supplementary Fig. 7.** ncWNT signaling pathway among rare subpopulations of BC.
**Supplementary Fig. 8.** The results for GiniClust on mouse retina datasets. We used the same marker genes with MarsGT and annotated the cell populations.
**Supplementary Fig. 9.** The eGRN of the structural constituent of eye lens pathway.
**Supplementary Fig. 10.** Heatmap of DEGs among all cell populations. The red box represents marker genes that have been reported by previous studies.
**Supplementary Fig. 11.** VEGF signaling pathway among major cell populations.
**Supplementary Fig. 12.** The dotplot of the accessibility of marker genes corresponding to enhancers.
**Supplementary Fig. 13.** The cell cluster results by Seurat, the circle means B cells which

are annotated by the curated marker genes.

**Supplementary Fig. 14.** The observed and extrapolated future states (arrows) after the knockout of MEF2C, NFIC, and SPI1 on the four subtypes of B cells.

**Supplementary Fig. 15.** The heatmap of DEGs expression in each cell type.

**Supplementary Fig. 16.** The heatmap of DEGs expression in three CD8+ cell types.

**Supplementary Fig. 17.** The introduction of the data information.

**Supplementary Fig. 18.** The regulatory relations of gene PRF1, GZMB, and IFNG.

**Supplementary Table S1**

**Supplementary Data (separate file)**

**Supplementary Data 1.** Datasets used in this paper for benchmarking and case study.

**Supplementary Data 2.** The evaluation of simulation datasets for benchmarking.

**Supplementary Data 3.** The evaluation of simulation datasets for the false positive test.

**Supplementary Data 4.** The evaluation of real datasets in different percentages of rare cell populations for benchmarking.

**Supplementary Data 5.** Grid optimization MarsGT on real bench sets.

**Supplementary Data 6.** The evaluation of real datasets for benchmarking.

**Supplementary Data 7.** Twenty repeated tests for robustness.

**Supplementary Data 8.** The cell-cell communication among BC types (case 1).

**Supplementary Data 9.** The enhancer-gene network with "the structural constituent of eye lens" pathway (case 1).

**Supplementary Data 10.** The enhancer-gene network in cell type 2 and cell type 10 (case 1).

**Supplementary Data 11.** The difference enhancer-gene network in cluster 2 and cluster 10 (case 1).

**Supplementary Data 12.** Top 10 DEGs in each cell type (case 1).

**Supplementary Data 13.** The cell-cell communication among major cell types (case 1).

**Supplementary Data 14.** The gene signature in three pathways (case 3).

**Supplementary Data 15.** The pathway enrichment score in each B cell type (case 2).

**Supplementary Data 16.** The enhancer-gene network in each B cell type (case 2).

**Supplementary Data 17.** The DEGs in clusters 1, 9, and 12 (case 3).

**Supplementary Data 18.** The marker gene signature of NKT and MAIT (case 3).

**Supplementary Data 19.** The enhancer-gene network in clusters 1, 9, and 12 (case 3).

**Supplementary Data 20.** The marker gene signature of exhausted and effector (case 3).

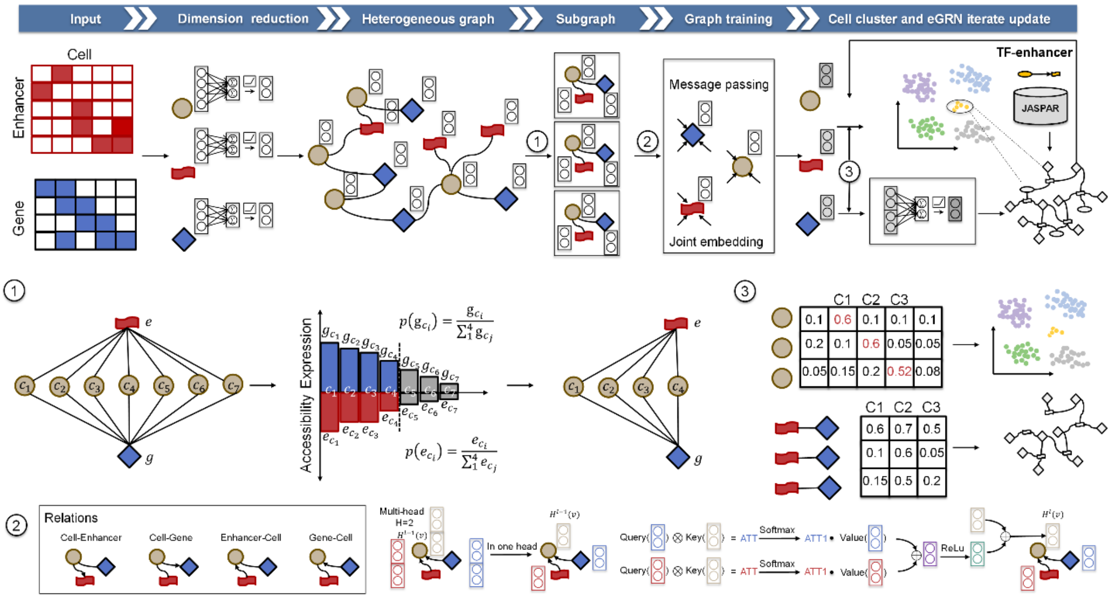


**Supplementary Fig. 1.** The detailed workflow of MarsGT for rare cell population identification.

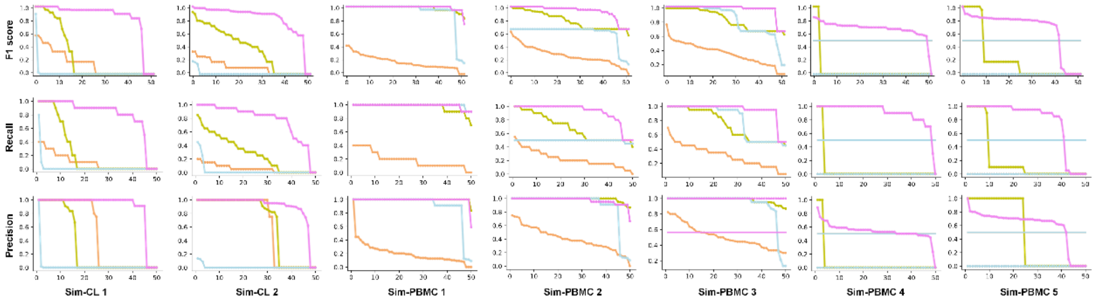


**Supplementary Fig. 2.** Performance comparison of rare cell identification on 350 simulated datasets in terms of F1 score, Precision, and Recall.

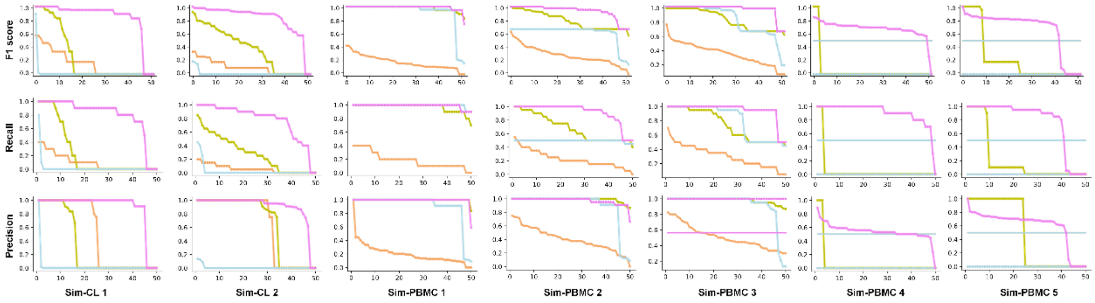


**Supplementary Fig. 3.** Performance comparison of the major cell and rare cell population simultaneously identification ability on 350 simulated datasets in terms of NMI, Purity, and Entropy.

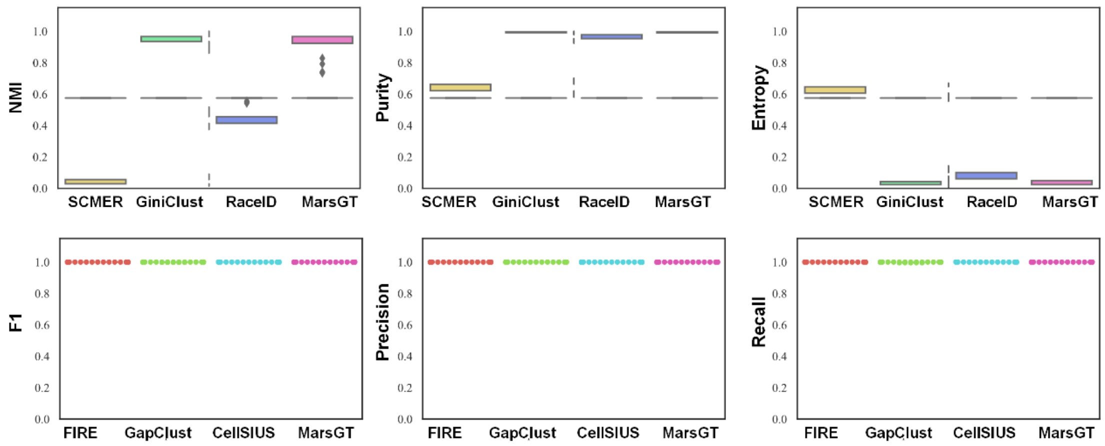


**Supplementary Fig. 4.** Performance comparison of the false positive rate on 50 simulated datasets in terms of F1 score, Precision, Recall, NMI, Purity, and Entropy.

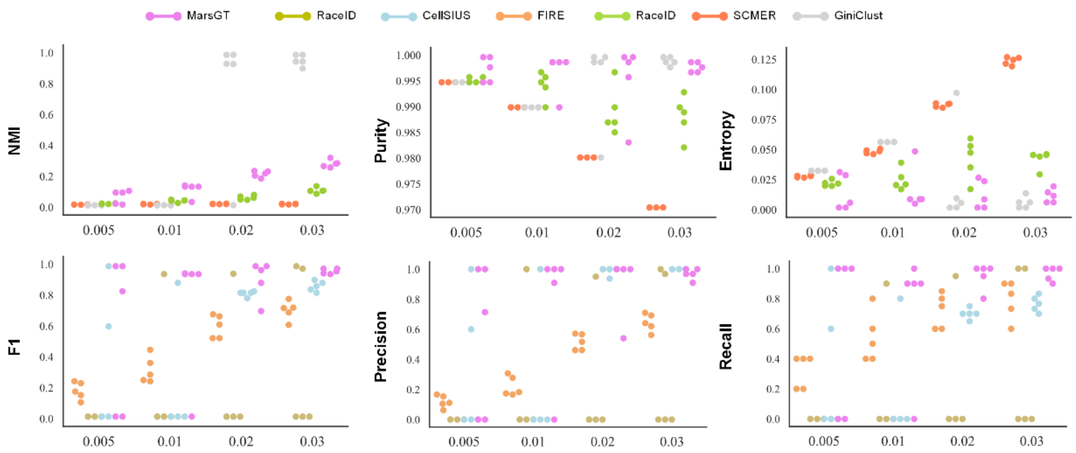


**Supplementary Fig. 5.** Performance comparison of the different proportion rare cell population identification ability on 5 simulated datasets in terms of F1 score, Precision, Recall, NMI, Purity, and Entropy.

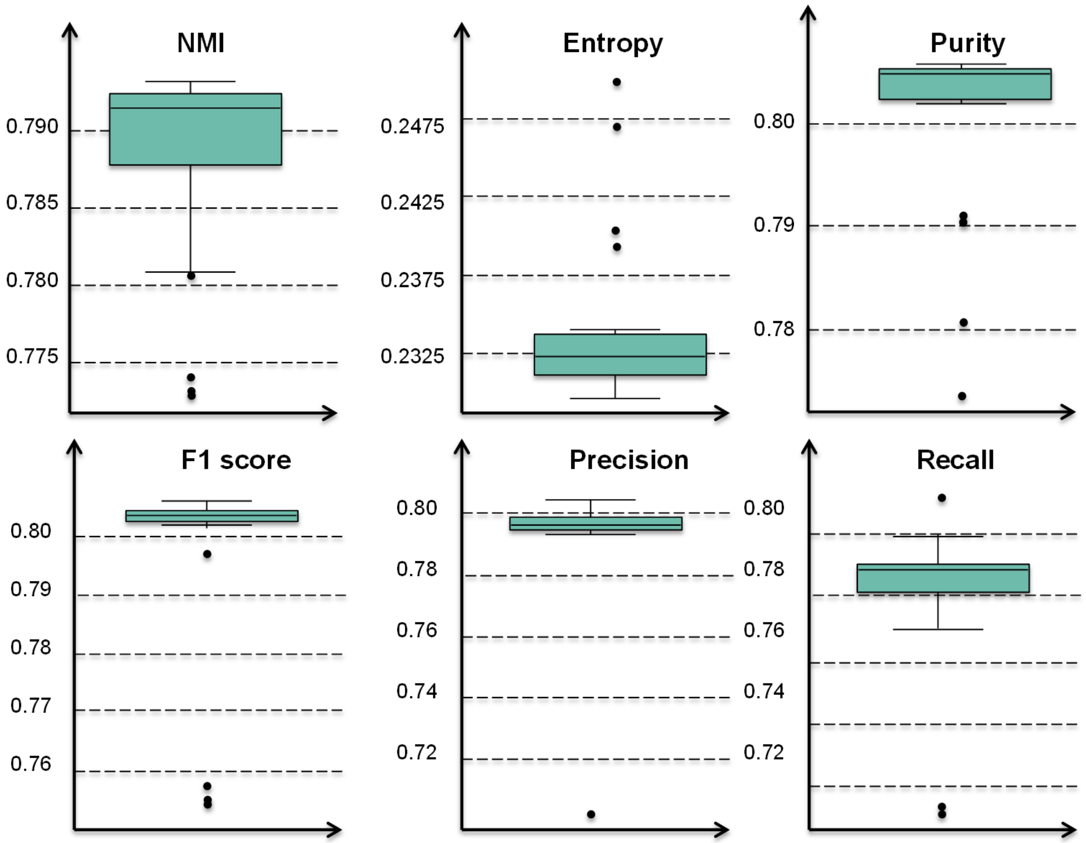


**Supplementary Fig. 6.** Robustness test. MarsGT runs 20 times on the independent test set.

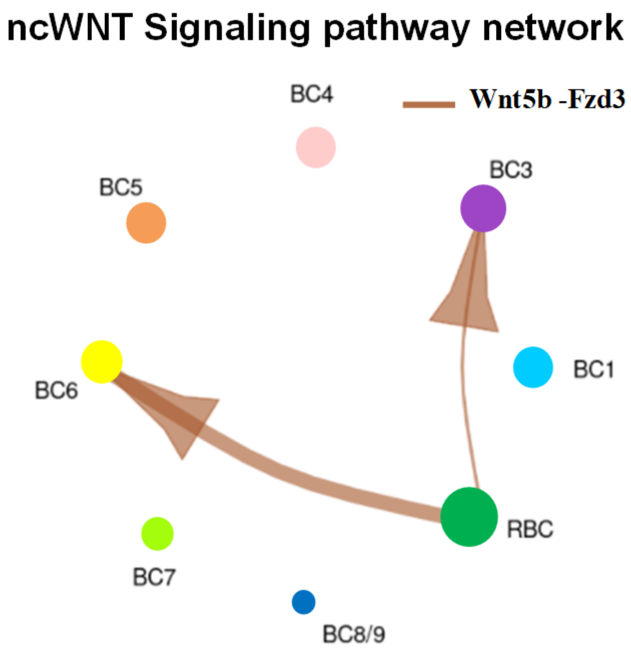


**Supplementary Fig. 7.** ncWNT signaling pathway among rare subpopulations of BC. A link between a filled circle (resource cluster with highly expressed ligand coding genes) and an unfilled circle (target cluster with highly expressed receptor coding genes) indicates the potential cell-cell communication of a signaling pathway. Circle colors represent different cell clusters, and the size represents the number of cells.

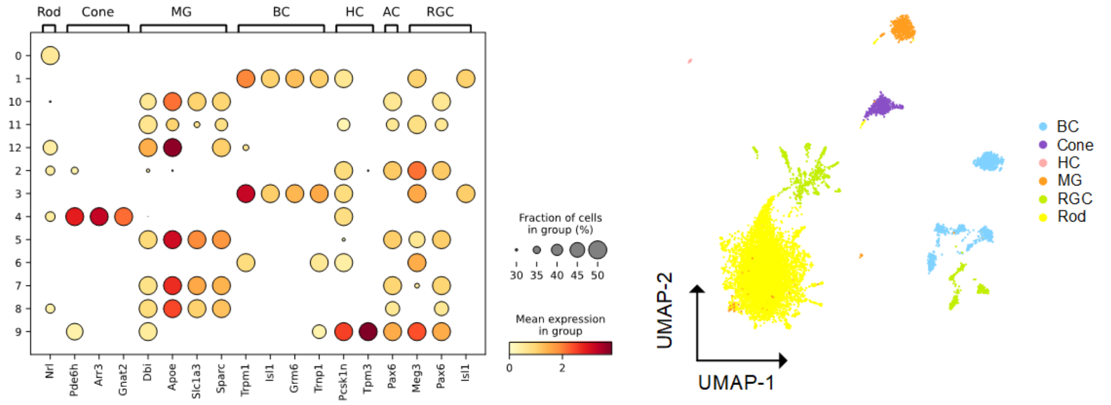


**Supplementary Fig. 8.** The results for GiniClust on mouse retina datasets. We used the same marker genes with MarsGT and annotated the cell populations.

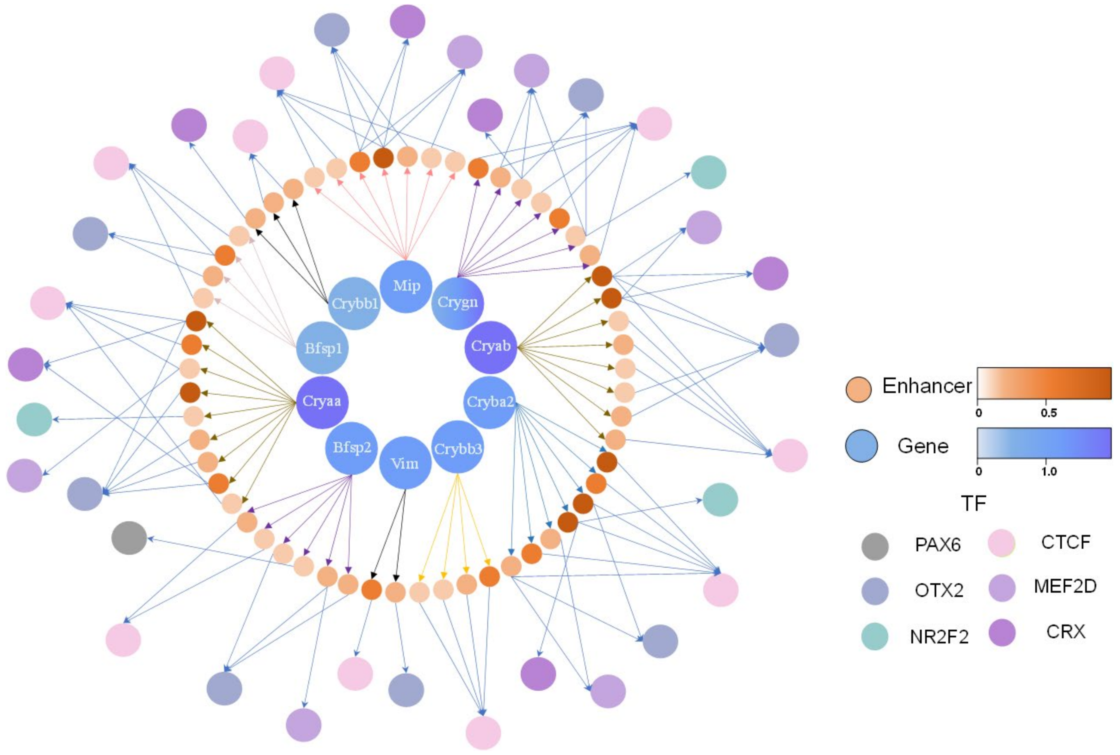


**Supplementary Fig. 9.** The eGRN of the structural constituent of eye lens pathway. The outer circles represent TFs. The inner circles represent genes in the structural constituent of eye lens pathway. The intermediate circles represent enhancers. The color of genes/enhancers represents the expression/accessibility.

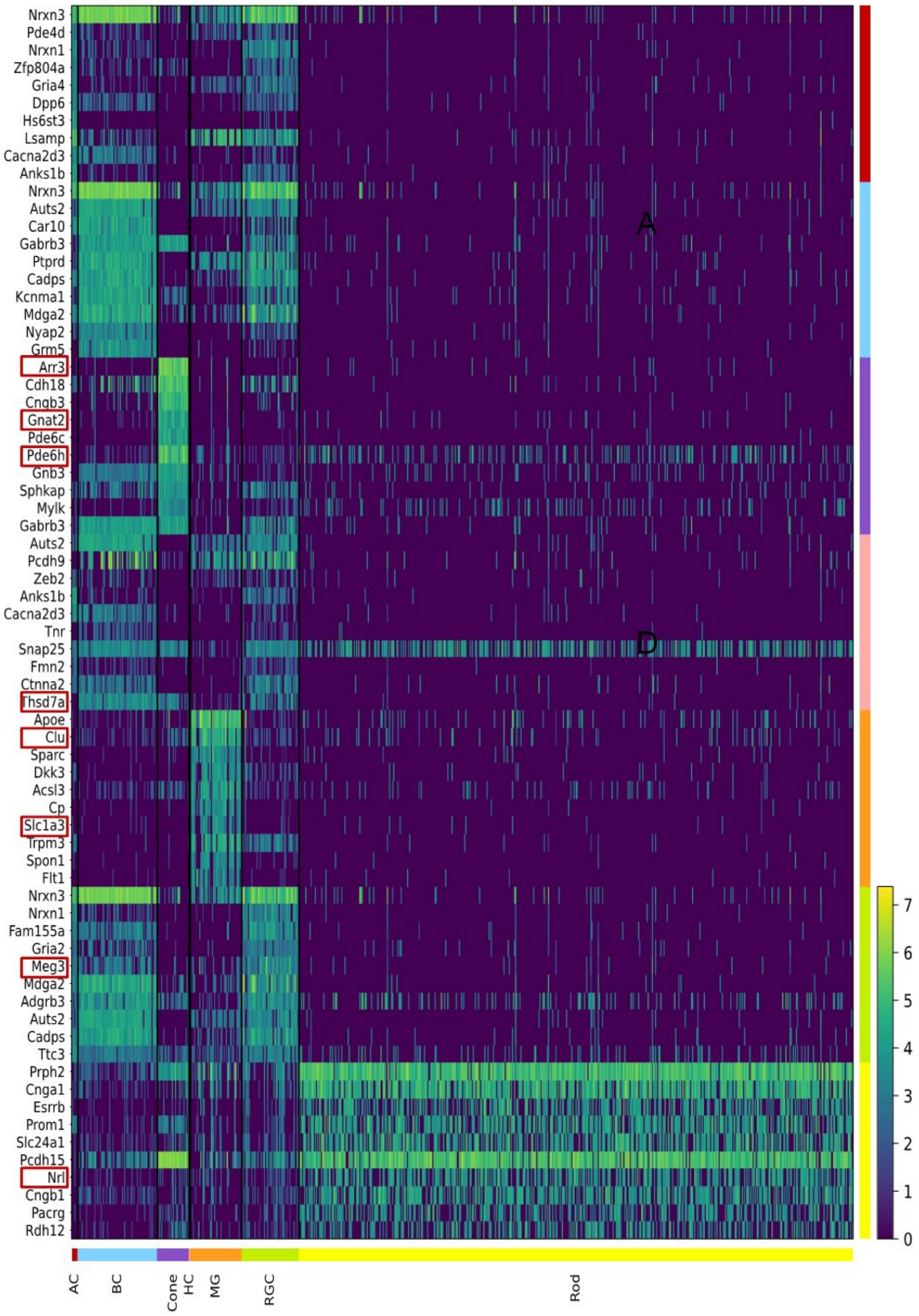


**Supplementary Fig. 10.** Heatmap of DEGs among all cell populations. The red box represents marker genes that have been reported by previous studies.

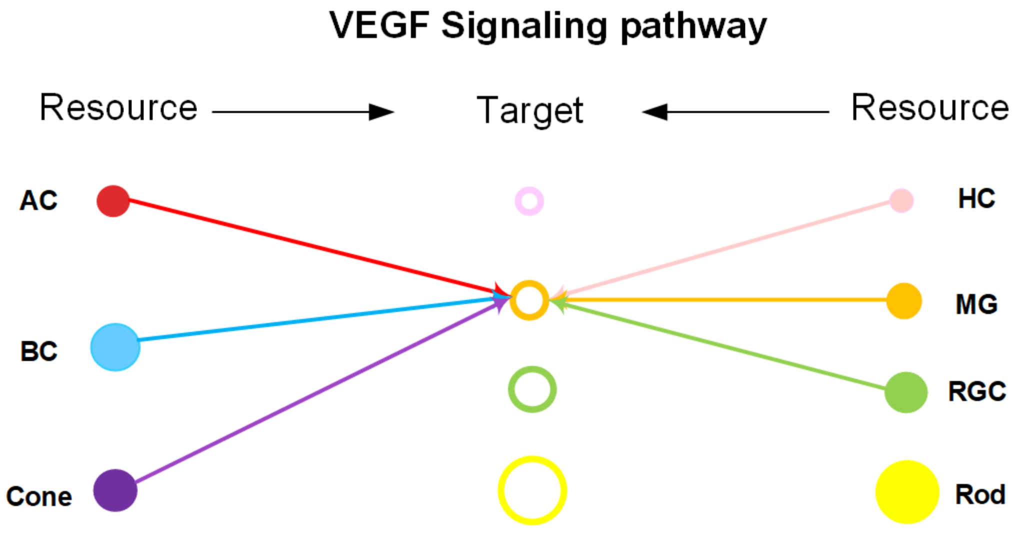


**Supplementary Fig. 11.** VEGF signaling pathway among major cell populations. A link between a filled circle (resource cluster with highly expressed ligand coding genes) and an unfilled circle (target cluster with highly expressed receptor coding genes) indicates the potential cell-cell communication of a signaling pathway. Circle colors represent different cell clusters, and the size represents the number of cells.

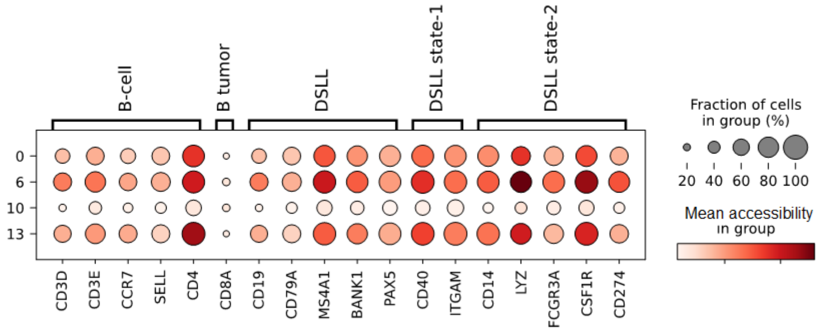


**Supplementary Fig. 12.** The dotplot of the accessibility of marker genes corresponding to enhancers. The size of the dot means the fraction of cells, and the color represents the accessibility in the group.

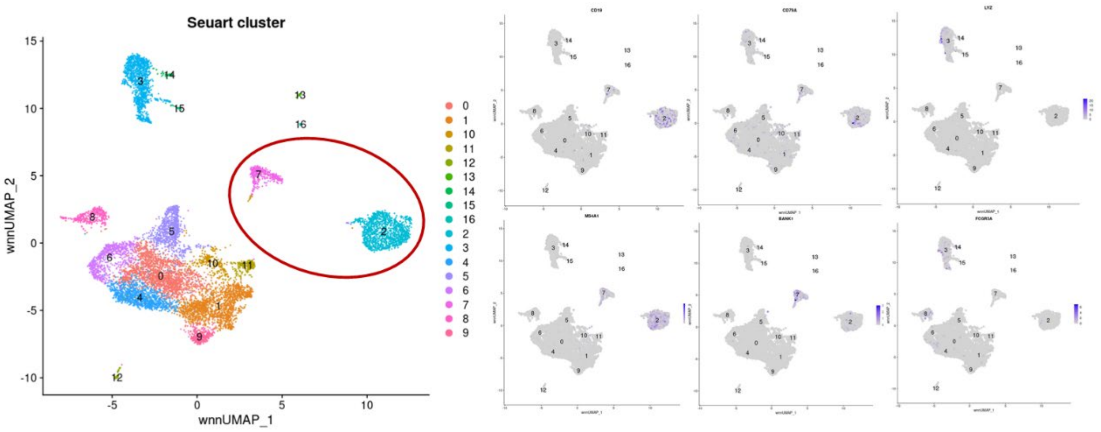


**Supplementary Fig. 13.** The cell cluster results by Seurat, the circle means B cells which are annotated by the curated marker genes.

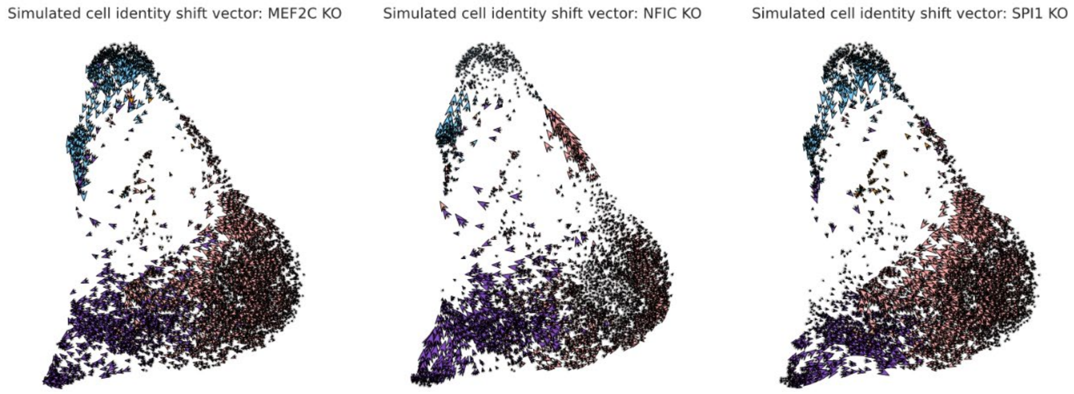


**Supplementary Fig. 14.** The observed and extrapolated future states (arrows) after the knockout of MEF2C, NFIC and SPI1 on the four subtypes of B cells. The color represents the cell clusters.

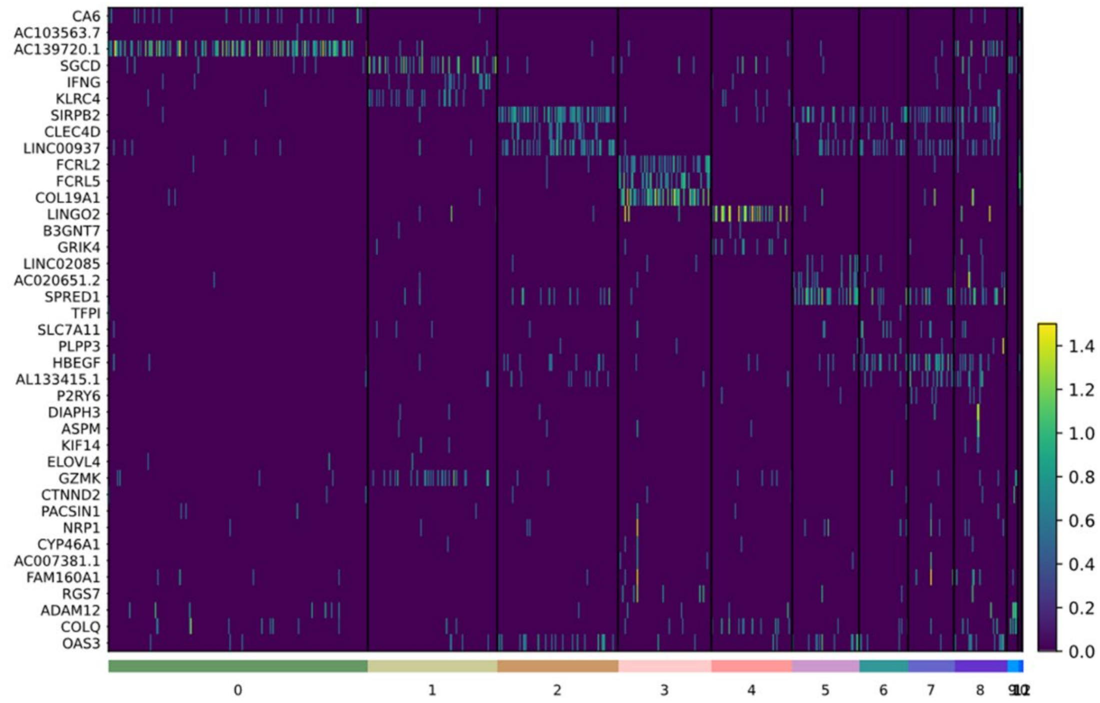


**Supplementary Fig. 15.** The heatmap of DEGs expression in each cell type.

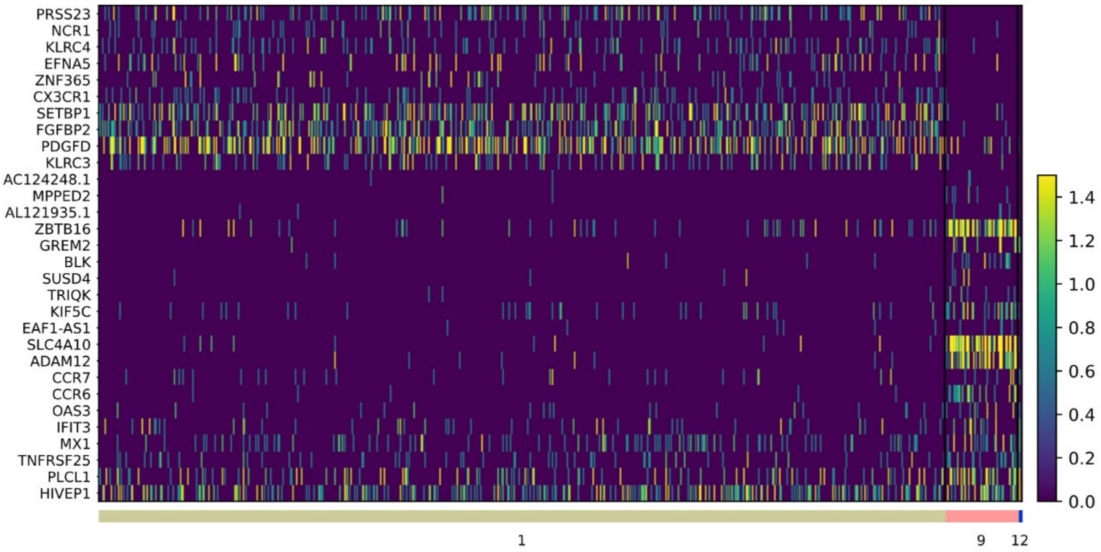


**Supplementary Fig. 16.** The heatmap of DEGs expression in three CD8+ cell types.

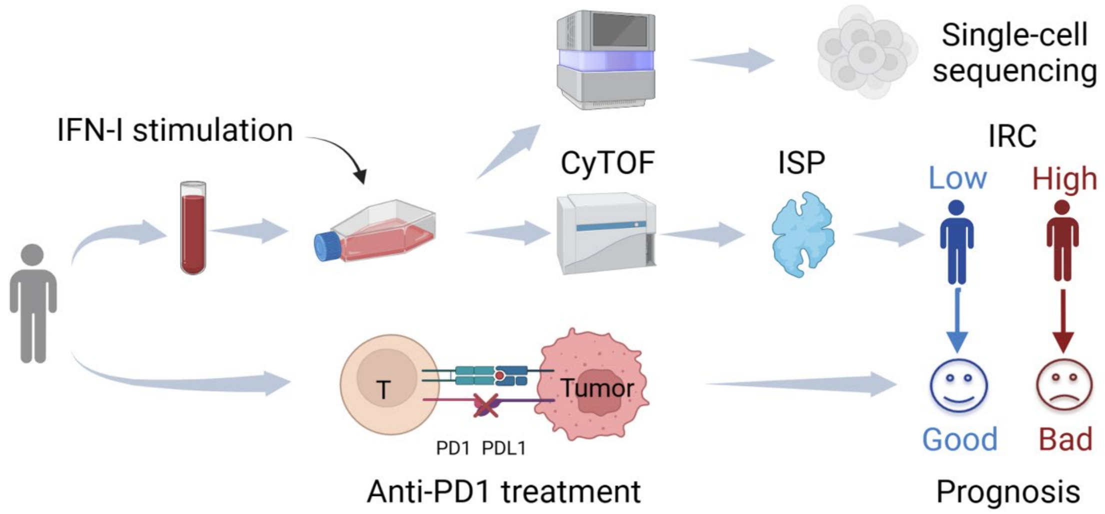


**Supplementary Fig. 17.** The introduction of the data information.

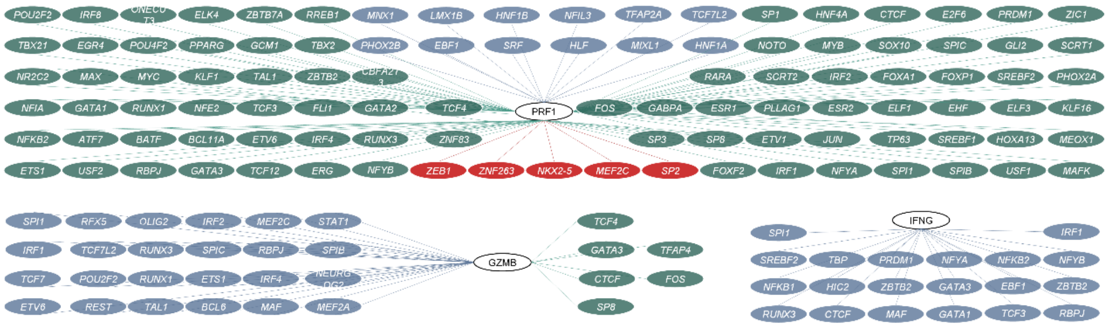


**Supplementary Fig. 18.** The regulatory relations of gene PRF1, GZMB, and IFNG. The red ellipse means the regulatory relation only exists in the high IRC, the blue ellipse means the regulatory relation only exists in the low IRC, and the green ellipse means the common regulatory relation between the high IRC and low IRC.

**Supplementary Table S1: Cell Line and PBMC simulated Datasets**

| Dataset Type | Common Cell Types | Rare Cell Types |
|---|---|---|
| Sim-CL 1 | PDX1, PDX2 (290 cells) | HeLa.S3 (10 cells) |
| Sim-CL 2 | PDX1, PDX2 (280 cells) | HeLa.S3 (10 cells), K562 (10 cells) |
| Sim-PBMC 1 | CD8+T (490 cells) | Plasma (10 cells) |
| Sim-PBMC 2 | CD4+T naïve (480 cells) | HSC (10 cells), Plasma (10 cells) |
| Sim-PBMC 3 | CD8+T (490 cells) | Erythroblast (10 cells) |
| Sim-PBMC 4 | CD8+T (480 cells) | Erythroblast (10 cells), HSC (10 cells) |
| Sim-PBMC 5 | CD8+T (480 cells) | Erythroblast (10 cells), Naive CD20+B (10 cells) |
| Sim-PBMC 6 | CD14+Mono (250 cells), CD8+ (250 cells) | - |
